# Histone H1 limits DNA methylation in *Neurospora crassa*

**DOI:** 10.1101/041285

**Authors:** Michael Seymour, Lexiang Ji, Alex M. Santos, Masayuki Kamei, Takahiko Sasaki, Evelina Y. Basenko, Robert J. Schmitz, Xiaoyu Zhang, Zachary A. Lewis

## Abstract

Histone H1 variants, known as linker histones, are essential chromatin components in higher eukaryotes, yet compared to the core histones relatively little is known about their in vivo functions. The filamentous fungus *Neurospora crassa* encodes a single H1 protein that is not essential for viability. To investigate the role of *N. crassa* H1, we constructed a functional FLAG-tagged H1 fusion protein and performed genomic and molecular analyses. Cell fractionation experiments showed that H1-FLAG is a chromatin binding protein. Chromatin-immunoprecipitation combined with sequencing (ChIP-seq) revealed that H1-3XFLAG is globally enriched throughout the genome with a subtle preference for promoters of expressed genes. In mammals, the stochiometery of H1 impacts nucleosome repeat length. To determine if H1 impacts nucleosome occupancy or nucleosome positioning in *N. crassa*, we performed Micrococcal nuclease digestion in wildtype and the Δ*hH1* strain followed by sequencing (MNase-seq). Deletion of *hH1* did not significantly impact nucleosome positioning or nucleosome occupancy. Analysis of DNA methylation by whole-genome bisulfite sequencing (MethylC-seq) revealed a modest but global increase in DNA methylation in the Δ*hH1* mutant. Together, these data suggest that H1 acts as a non-specific chromatin binding protein that can limit accessibility of the DNA methylation machinery in *N. crassa*.

## INTRODUCTION

In eukaryotes, packaging of genomic DNA into chromatin is essential for genome function. The most basic unit of the chromatin fiber is the nucleosome core particle (NCP), made up of ~146 bp of DNA wrapped around a core of histones H3, H4, H2A and H2B (Kornberg and Thomas 1974; Olins and Olins 1974; Luger *et al.* 1997). In addition to the core histones many organisms encode one or more H1 proteins, also known as linker histones. H1 proteins are evolutionarily unrelated to the core histones and are characterized by a central winged helix domain, or globular domain, flanked by unstructured N-and C-termini (Cerf *et al.* 1993; Ramakrishnan *et al.* 1993; Kasinsky *et al.* 2001). Early studies showed that animal H1 proteins bind outside of the NCP (Baldwin *et al.* 1975; Shaw *et al.* 1976) and can protect an additional 20bp of DNA from nuclease digestion (Whitlock and Simpson 1976; Noll and Kornberg 1977). Subsequent studies revealed that H1 binds near the NCP dyad axis and can interact with DNA as it enters and exits the NCP (recently reviewed in Bednar *et al.* 2015). Although the interactions between H1 and the NCP have been extensively investigated, H1’s roles in the cell remain poorly understood.

In vivo studies of mammalian H1 are complicated by the existence of 11 H1 variants that appear to be partially redundant (Pan and Fan 2015). Deletion of single H1 variants failed to produce significant phenotypes in mice (Fan *et al.* 2001), but mice lacking three H1 variants are inviable (Fan *et al.* 2003) and triple-knockout embryonic stem cells (ESCs) are unable to differentiate (Zhang *et al.* 2012a). These and other data suggest that animal H1 variants cooperate to perform critical functions, influencing gene regulation (Fan *et al.* 2005b; Li *et al.* 2012b; Zhang *et al.* 2012b), establishment and/or maintenance of chromatin modification patterns (Li *et al.* 2012b; Zhang *et al.* 2012a; Yang *et al.* 2013; Lu *et al.* 2013) and formation of higher order chromatin structures (Fan *et al.* 2005b; Geeven *et al.* 2015).

Less is known about the functions of H1 in other groups of organisms, but genetic studies have been carried out in a handful of microbial model systems. H1 is not essential for viability in the single-celled *Saccharomyces cerevisiae* (Patterton *et al.* 1998) or *Tetrahymena thermophila* (Shen *et al.* 1995). Similarly, H1-deficient mutants are viable in several filamentous fungi including *Neurospora crassa* (Folco *et al.* 2003), *Aspergillus nidulans* (Ramón *et al.* 2000), and *Ascobolus immersus* (Barra *et al.* 2000). The yeast H1 homolog Hho1p suppresses homologous recombination (Downs *et al.* 2003; Li *et al.* 2008), impacts ribosomal RNA processing (Levy *et al.* 2008), and influences chromatin compaction during stationary phase (Schäfer *et al.* 2008). In *T. thermophila*, H1 is required for normal chromatin compaction in macronuclei and influences expression of a small number of genes (Shen *et al.* 1995; SHEN and GOROVSKY 1996). It is important to note that both yeast Hho1p and *T. thermophila* H1 have atypical protein structures. The yeast protein contains two globular domains, whereas the *T. thermophila* protein lacks a globular domain altogether. Thus, it is not clear if these proteins are functionally analogous to H1 in other organisms.

The filamentous fungi *N. crassa*, *A. nidulans*, and *A. immersus* encode H1 proteins with a canonical tripartite structure, raising the possibility that these genetic systems can be used to gain insights into H1 function in plants and animals. In *N. crassa*, an *hH1*-deficient strain displayed reduced growth and H1 was required for repression of the *cfp* gene in the presence of ethanol (Folco *et al.* 2003). In *A. immersus*, H1 gene silencing led to increased nuclease accessibility and increased DNA methylation (Barra *et al.* 2000). In contrast, deletion of *hhoA* in *A. nidulans* failed to produce a phenotype (Ramón *et al.* 2000). In general, the functions of H1 in filamentous fungi remain poorly understood. Moreover, it is not clear if H1 plays similar roles in fungal and animal cells. In the present study, we utilized molecular and genomic approaches to investigate the functions of H1 in the model fungus *N. crassa*. We confirmed that *N. crassa* H1 is a chromatin component in vivo, and we found that H1 is not a major determinant of nucleosome positioning, nucleosome repeat length, or nucleosome occupancy. We report that an H1 fusion protein exhibits enhanced enrichment at nucleosome free regions in a ChIP-seq assay and that loss of H1 causes a global increase in DNA methylation.

## >MATERIALS AND METHODS

### Strains, Growth Media, and Molecular Analyses

All Neurospora strains used in this study are listed in **Supplementary File 1**. Knockout strains of hH1 were generated by the Neurospora gene knockout consortium (Colot *et al.* 2006) and obtained from the Fungal Genetics Stock Center (McCluskey *et al.* 2010). Neurospora cultures were grown at 32° in Vogel’s minimal medium (VMM) + 1.5% sucrose (Davis *et al.* 1970). Crosses were performed on modified synthetic cross medium at 25° (Davis *et al.* 1970). For plating assays, Neurospora conidia were plated on VMM with 2.0% sorbose, 0.5% fructose, and 0.5% glucose. When relevant, plates included 200 *μ*g/mL hygromycin or 400 *μ*g/mL basta (Pall 1993). Neurospora transformation (Margolin *et al.* 1997), DNA isolation (Pomraning *et al.* 2009), protein isolation and Western blotting (Honda and Selker 2008) were performed as previously described. 3χ-FLAG knock-in constructs were made by introduction of linear DNA fragments constructed by overlapping PCR using described plasmid vectors (Honda and Selker 2009) and *N. crassa* fragments generated using the following primers: H1 CDS FP 5′- GAG GTC GAC GGT ATC GAT AAG CTT AT ATC CAC CGA CAA CAT GTT CGA CTC - 3′; H1 CDS RP 5′ - CCT CCG CCT CCG CCT CCG CCG CCT CCG CCT GCC TTC TCG GCA GCG GGC TC −3′; H1 UTR FP 5′ - TGC TAT ACG AAG TTA TGG ATC CGA GCT CGA CTC GTT CCT TTG GGA TGA T - 3′; H1 UTR RP 5′ - ACC GCG GTG GCG GCC GCT CTA GAA CTA GTT CAT CAA ACC AAA TTC TCG G - 3′. To separate soluble nuclear proteins from the chromatin fraction, cultures were grown overnight and cells were collected, ground in liquid Nitrogen, and resuspended in 1 mL of low salt extraction buffer (50mM HEPES-KOH pH 7.5, 150mM NaCl, 2mM EDTA, plus protease inhibitor tablets (cat # 11836153001; Roche, Indianapolis, IN)). Extracts were centrifuged at 14,000 rpm and the supernatant containing soluble proteins was saved. The pellet was resuspended in 1mL of high salt extraction buffer (50mM HEPES-KOH pH 7.5, 600mM NaCl, 2mM EDTA, plus protease inhibitor tablets (cat # 11836153001; Roche, Indianapolis, IN)) and subjected to sonication. Extracts were centrifuged at 14,000 rpm in a microfuge and the supernatant was saved as the chromatin fraction. Both fractions were analyzed by Western blotting using anti-FLAG antibodies (cat # F1804; Sigma-Aldrich) and anti-H3 antibodies (cat # 06-755 Millipore).

### DNA Sequencing and Data Analysis

#### ChIP-seq

For chromatin immunoprecipitation (ChIP) experiments, 5 ×10^6^ conidia/ml were inoculated into 50 mL of VMM and incubated at 32° for 5 hours. Cells were harvested and ChIP was performed as previously described (Sasaki *et al.* 2014) using anti-FLAG antibodies (cat # F1804; Sigma-Aldrich) or antibodies to the unphosphorylated C-terminal repeat of RNA polymerase II (8WG16; cat# MMS-126R; Covance). Two biological replicates were performed for each experiment. Libraries were prepared using the TruSeq ChIP sample prep kit (Illumina, cat # IP-202-1012) according to manufacture instructions with the following modification. Library amplification was performed using only 4 cycles of PCR to reduce biased enrichment of GC-rich DNA (Ji *et al.* 2014). Libraries were sequenced at the University of Georgia Genomics Facility on an Illumina NextSeq 500 instrument. Reads were aligned to version 12 of the *N. crassa* genome (Refseq Accession # GCF_000182925.2; Galagan *et al.* (2003)) using the Burrows-Wheeler Alingner (BWA version 0.7.10) (Li and Durbin 2009). To determine if H1-FLAG was enriched over background, coverage was normalized to mitochondrial DNA as follows. We used BEDtools (version 2.25.0) ‘coverage’ to calculate read coverage for 1000bp windows across the genome (Quinlan and Hall 2010). We then used BEDtools ‘map’ to calculate the median coverage for mitochondrial DNA. The coverage for each 1000 base pair window was then divided by the median coverage for mitochondrial DNA. As a positive control, data from a previously published ChIP experiment for methylated lysine-9 of H3 was analyzed (Accession #SRX550120; Sasaki *et al.* 2014).

The Hypergeometric Optimization of Motif EnRichment (HOMER verson 4.7.2) (Heinz *et al.* 2010) software package (annotatePeaks.pl module) was used to generate metaplots and heatmaps of enrichment data (using the-hist and-ghist option, respectively). We first created a custom HOMER genome annotation for Neurospora using a fasta file and a GTF file (**Supplementary File 2**) containing the version 12 genome assemblies and annotations, respectively (Galagan *et al.* 2003). All plots were centered on transcriptional start sites or transcriptional termination sites and a window size of 10 bp was specified for all histograms (-hist 10). HOMER was also used to construct metaplots of expression-ranked gene groups using the-list option.

#### MNase digestion

For Micrococcal nuclease (MNase) experiments, 5 ×10^6^ conidia/mL were inoculated into 50 mL of VMM and incubated at 32° for 5 hours. The cell suspension was transferred to a 50mL conical and centrifuged at 1000 × g for five minutes to pellet germinated conidia. Cell pellets were washed with 10mL Phosphate Buffered Saline (PBS) (Sambrook *et al.* 1989) and then resuspended in 10mL of PBS containing 1% formaldahyde. The cell suspension was transferred to a 125mL flask and incubated with gentle shaking for 30 minutes at room temperature before the crosslinking agent was quenched by addition of 500*μ*L 2.5M Glycine. The cell suspension was transferred to a 50mL conical tube and germinating conidia were pelleted by centrifugation for 5 minutes at 1000 × g. Cells were washed once in 40mL of PBS and resuspended in 1mL of ice-cold PBS. Cells were pelleted by centrifugation for 5 minutes at 5000 × g, and each cell pellet was resuspended in NPS buffer with Calcium Chloride (50mM HEPES-KOH, pH 7.5, 140mM NaCl, 5mM MgCl_2_, 1mM CaCl_2_, 1% Triton-χ 100, 0.1% Deoxycholate, 0.5mM Spermidine, 1mM PMSF plus Roche EDTA-free protease inhibitor tablets [catalog # 05892791001]). Cells were then lysed by gentle sonication using a Branson sonicator with micro tip (Output 2.0, Duty Cycle 80%; 30 1-second pulses). The chromatin fraction was pelleted by centrifugation for 5 minutes at 14,000 × g. The supernatent was discarded and each pellet was resuspended in 1 mL of NPS buffer and transferred to a 15mL conical tube. NPS buffer with Calcium Chloride was added to raise the volume to 6 mL and the chromatin sample was mixed by pipetting. 700 *μ*L aliquots were transferred to 1.5 mL tubes and 2 units of Micrococcal nuclease (cat #2910A; Clonetech) were added to each tube. Individual samples were incubated at 37° for 5, 10, 20, 40, 60, 90, or 120 minutes. MNase digestions were stopped by addition of 15*μ*L 0.5M EDTA and 25 *μ*L 4M NaCl and samples were incubated overnight at 65° to reverse cross-links. 6*μ*L RNase A (10mg/ml; Fisher Scientific, cat # BP2529250) was added and samples were incubated for 2 hours at 50°. 6*μ*L of 10% SDS and 10*μ*L Proteinase K (10mg/ml; Fisher Scientific, cat # BP1700) was then added and samples were incubated for 2 hours at 65°. The digested DNA was isolated by phenol-chloroform extraction and precipitated overnight at −20° in ethanol and sodium acetate (Sambrook *et al.* 1989). Digested DNA was resolved by gel electrophoresis to confirm that digestion was successful.

#### MNase-seq

We constructed sequencing libraries from mono-nucleosomes generated by 20 or 60 minute MNase digestion. We first performed a gel extraction (Qiagen, cat #28706) of mononucle-somal DNA (~150bp) and constructed libraries using an Illumina TruSeq Sample preparation kit (Illumina) according to manufacture instructions. 50bp paired-end sequencing reads were generated on a Illumina HiSeq 2500 instrument at the Oregon State University genomics core facility. Due to a technical problem during the sequencing run, only 44 bp of sequence were obtained for the read 2 sequence. Sequences were mapped to the *N. crassa* version 12 genome assembly (Galagan *et al.* 2003) using bowtie2 (version 2.2.3) (Langmead and Salzberg 2012). To analyze the size distributions in wildtype and the hH1 strain, the Picard software package (http://broadinstitute.github.io/picard) was used to remove duplicate reads and determine insert size metrics (using ‘CollectInsertSizeMetrics’). HOMER was used to create metaplots of MNase data as described above. In all cases, a window size of 10bp was used (-hist 10). Metaplots depict only plus strand reads (using the ‘-strand +’ option) and thus peaks indicate the left edge of nucleosomes.

#### RNA-seq

For RNA-seq experiments, 5×10^6^ conidia/mL were inoculated into 50mL of VMM containing 2% glucose and grown for 5 hours at 32°. RNA isolation was performed as described (Bell-pedersen *et al.* 1996; Schwerdtfeger and Linden 2001) and strand-specific RNA-seq libraries were prepared from 5 *μ*g total RNA. Ribosomal RNAs were depleted using the yeast Ribo-zero kit (cat # MRZY1324 Epicentre) and RNA libraries were generated with the Illumina Stranded RNA-seq kit (cat # RS-122-2101). Reads were aligned to version 12 of the *N. crassa* genome sequence using TopHat (Trapnell *et al.* 2009) and expression levels (FPKM) were determined using Cufflinks (Trapnell *et al.* 2012).

#### MethylC-seq

For DNA methylation analysis, conidia were inoculated into 5 mL of VMM and cultures were grown for 48 hours. Genomic DNA was isolated using described procedures (Pomraning *et al.* 2009). MethylC-seq libraries were prepared as previously described (Urich *et al.* 2015). Illumina sequencing was performed at the University of Georgia Genomics Facility using an Illumina NextSeq 500 instrument. Sequencing reads were trimmed for adapters, preprocessed to remove low quality reads and aligned to the N. crassa version 12 genome assembly (Galagan *et al.* 2003) as described in Schmitz *et al.* (2013). Mitochondria sequence (which is fully unmethylated) was used as a control to calculate the sodium bisulfite reaction non-conversion rate of unmodified cytosines. Only cytosine sites with a minimum coverage (set as 3) were allowed for subsequent analysis. Binomial test coupled with Benjamini-Hochberg correction was adopted to determine the methylation status of each cytosine. Identification of DMRs (Differentially Methylated Regions) was performed as described (Schultz *et al.* 2015). Methylated regions in wild type were defined previously (Basenko *et al.* 2016). For metaplots, both upstream and downstream regions were divided into 20 bins each of 50bp in length for a total 1kb in each direction. Methylated regions were separated every 5%, for a total of 20 bins. Weighted methylation levels were computed for each bin as described previously (Schultz *et al.* 2012).

### Data Availability

All strains are listed in Supplementary File 1 and available upon request. All sequencing data have been deposited into the NCBI SRA/GEO databases. ChIP-seq, MNase-Seq, and RNA-seq data generated during this study have been deposited under accession #GSE78157 (reviewer link: http://www.ncbi.nlm.nih.gov/geo/query/acc.cgi?token=uvqzukkwhtgxfof&acc=GSE78157). Control data from a previously published ChIP-seq experiment for methylation of H3 lysine-9 was deposited under accession #SRX550120 (Sasaki *et al.* 2014). MethylC-seq data are deposited under accession #GSE76982 (this study; reviewer link: http://www.ncbi.nlm.nih.gov/geo/query/acc.cgi?token=edunyawudnqntwr&acc=GSE76982) and #GSE70518 (Basenko *et al.* 2016).

## RESULTS

### Construction of an H1-3XFLAG fusion protein

To investigate the role of H1 in N. crassa cells, we constructed an epitope-tagged version of the protein by introducing coding sequence for a 3χ-FLAG tag at the 3′ end of the native *hH1* locus (Figure 1A). Primary transformants were crossed to obtain a homokaryon that was analyzed further. To confirm that the H1-3XFLAG fusion protein is functional, we first compared the growth rate of the *hH1∷hH1-3xflag-hph* strain to wildtype and to an *hH1* deletion strain obtained from the Neurospora gene knockout consortium (Colot *et al.* 2006). The Δ*hH1* strain displayed a reduced growth rate, as reported previously for an H1 loss-of-function allele generated by repeat-induced point mutation (Folco *et al.* 2003). The *hH1∷hH1-3xflag-hph* grew similar to wildtype (Figure 1B and C). We also asked if the H1-3XFLAG protein associates with chromatin. We isolated soluble and chromatin-containing fractions (see Materials and Methods) and performed Western blot analyses using anti-FLAG and anti-H3 antibodies. Western blot probed with an anti-FLAG antibody revealed a single band. The apparent molecular weight was larger than expected based on amino acid sequence prediction, but the apparent size was consistent with previous analysis of *N. crassa* purified by extraction with perchloric acid (Folco *et al.* 2003). We detected H1-3XFLAG in both soluble and chromatin fractions with higher levels of H1-3XFLAG observed in the chromatin fraction (Figure 1D). As expected, H3 was exclusively detected in the chromatin fraction. Together, these data demonstrate that the H1-3XFLAG construct is functional and that *N. crassa* H1 is a component of chromatin.

**Figure 1.**
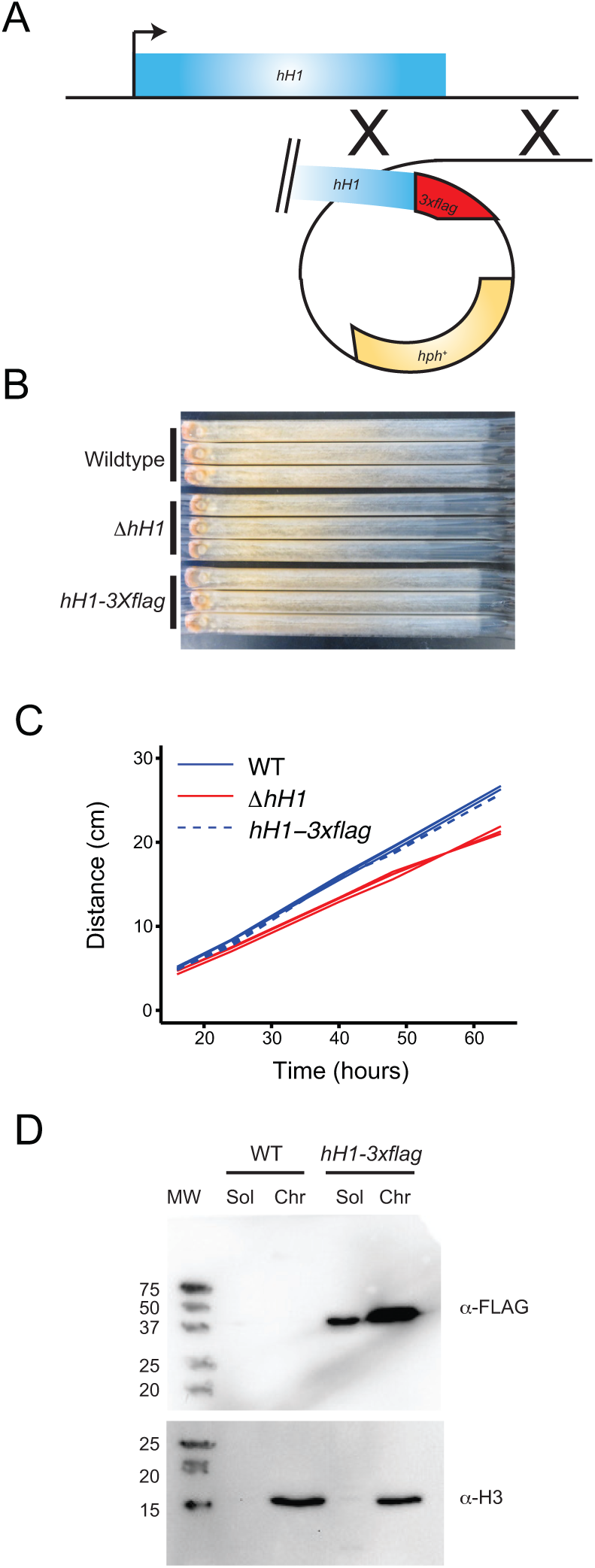
H1-3XFLAG is functional and binds to chromatin.(A) A cartoon illustrating the strategy for introducing a *3x-flag* sequence into the 3’ end of the native *hH1* locus by homologous recombination. (B) The linear growth rate of the indicated strains was measured using ‘race tubes’ for three replicates of each strain. The direction of growth is left to right. (C) Quantification of the linear growth rate data from panel B as distance (y-axis; cm) versus time (x-axis; hours). (D) The soluble (Sol) and the chromatin fractions (Chr) were isolated from wildtype and the *hH1-3xflag* strain and both fractions were analyzed by Western blotting using anti-FLAG and anti-H3 antibodies, as indicated. MW indicates the positions and sizes in kiloDaltons of a prestained protein ladder.

### H1 is moderately enriched in promoters and depleted from coding sequences of actively expressed genes

To determine the genome wide distribution of H1-3XFLAG, we performed chromatin-immunoprecipitation followed by sequencing (ChIP-seq). Inspection of the H1-3XFLAG enrichment data in a genome browser revealed a relatively uniform distribution for all seven chromosomes. Given that high levels of H1-3XFLAG were detected in the chromatin fraction by Western blotting, we reasoned that the uniform enrichment pattern observed for H1-3XFLAG might reflect global binding of H1 across the genome. To determine if this was the case, we normalized read counts obtained for each 1000 base pair window in the nuclear genome to the median read count obtained for all 1000 base pair windows covering the mitochondrial genome. This allowed us to calculate enrichment over background because mitochondrial DNA should not be enriched by immunoprecipitation of a nuclear protein. Data for all 1000 base pair windows on Linkage Group VII demonstrate that H1 is a general chromatin architectural protein in *N. crassa* ( Figure 2). ChIP-seq experiments using anti-FLAG antibodies led to global enrichment of chromosomal DNA from the *hH1-3xflag* strain but not from a wildtype negative control strain. As a positive control, we normalized read counts obtained in a previously published ChIP-seq experiment performed with antibodies to H3 methylated at lysine-9 (Sasaki *et al.* 2014). This confirmed that normalization to mitochondrial DNA is an effective method for quantifying enrichment over background.

**Figure 2.**
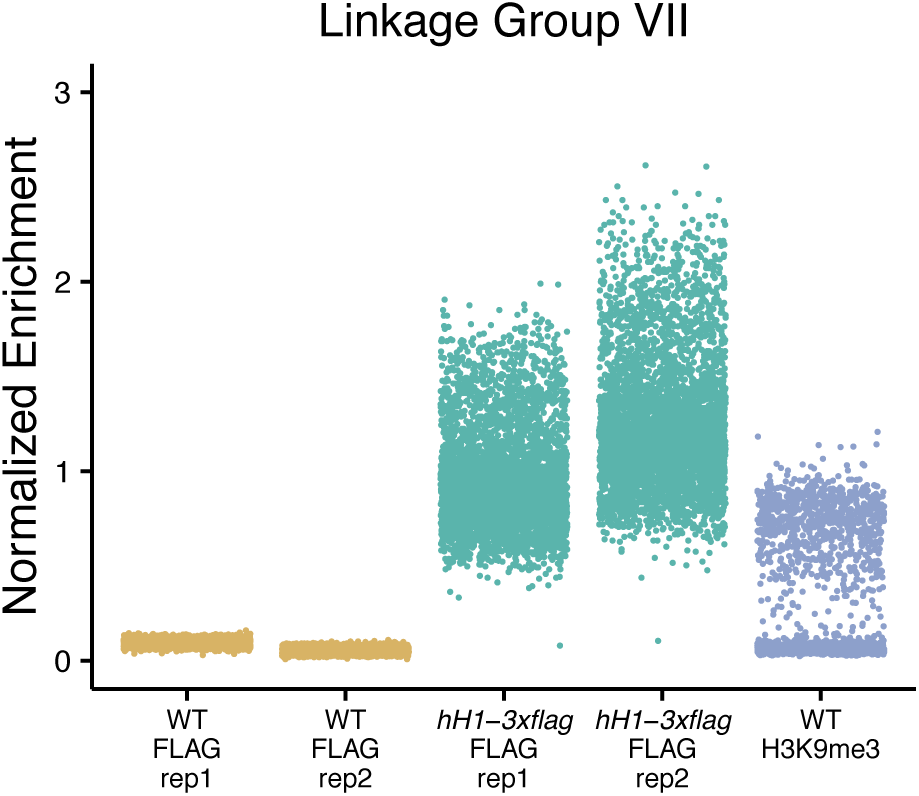
H1-3XFLAG is globally enriched throughout the *N. crassa* genome. Normalized read coverage is shown for all 1000 base pair windows across *N. crassa* LGVII for anti-FLAG ChIP-seq experiments performed with a wildtype negative control strain (no FLAG protein) and the *hH1-3xflag* strain (two replicates for each strain are shown). Normalized read coverage is also plotted for an anti-H3K9me3 ChIP-seq experiment performed with wildtype. Each spot depicts the read count for a single 1000bp window normalized to the median enrichment obtained for mitochondrial DNA windows.

We did not detect prominent peaks that are typical of transcription factors or previously analyzed histone modifications such as H3K4me2 or H3K9me3 (Lewis *et al.* 2009). However, we did detect a subtle enrichment of H1 in the promoters of many genes along with a corresponding depletion of H1-3XFLAG within many coding sequences. To determine if this pattern occurred broadly across the *N. crassa* chromatin landscape, we created metaplots to analyze the average H1 distribution across the transcriptional start sites (TSS) or transcriptional termination sites (TTS) of all *N. crassa* genes (Figure 3A).

**Figure 3.**
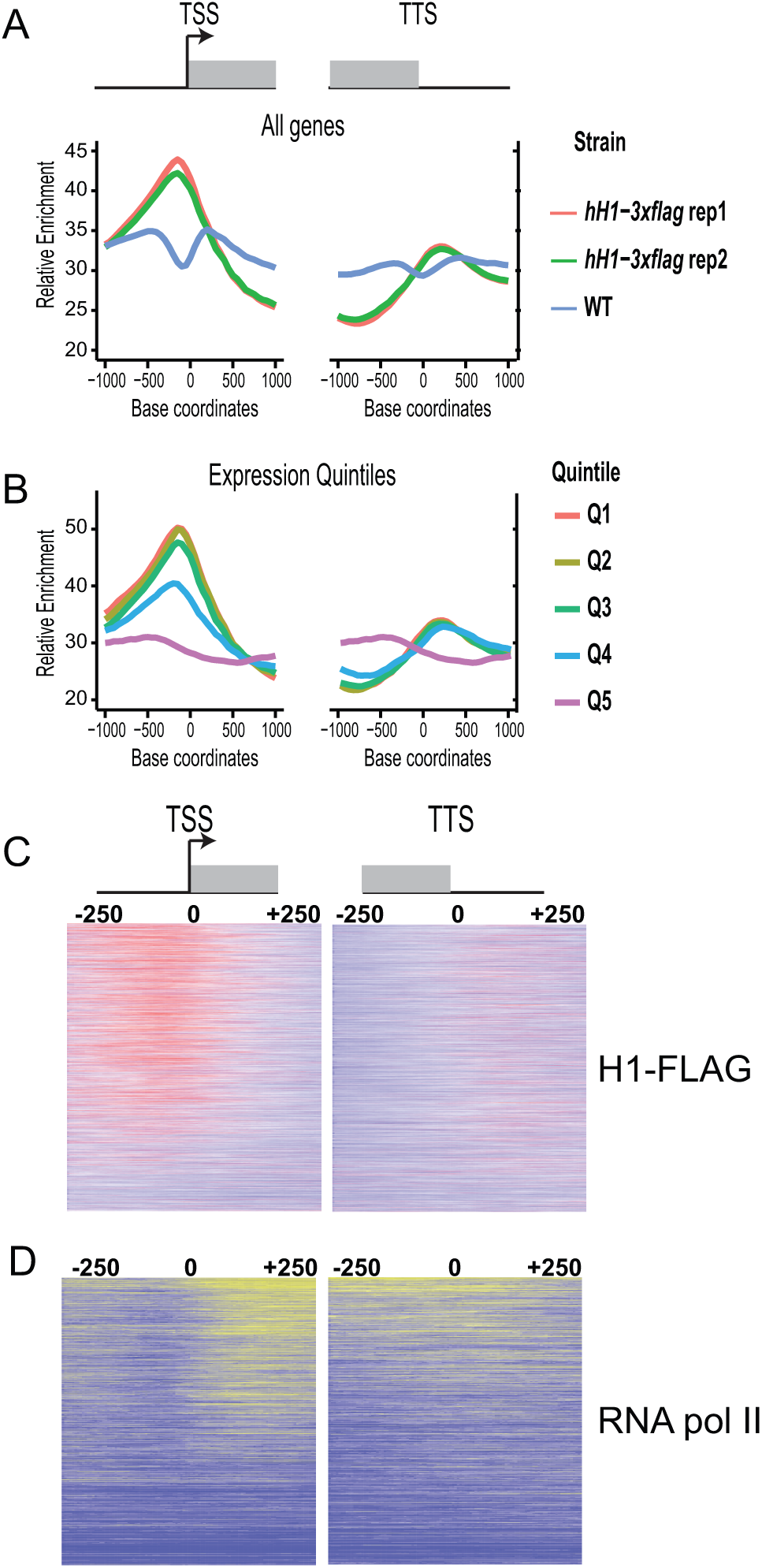
H1-3XFLAG is depleted from gene bodies and modestly enriched in promoters of expressed genes. (A) Metaplots depict the average ChIP-seq enrichment pattern across all *N. crassa* genes for two replicate FLAG ChIP-seq experiments performed with the *hH1-3xflag-hph*^*+*^ strain and a wildtype negative control strain (no FLAG-tagged protein). Metaplots are centered on the the transcrip-tional start (TSS; left) or the transcriptional termination site (TTS; right). (B) All *N. crassa* genes were ranked by expression level and split into quintile groups. Quintile 1 (Q1) corresponds to genes with the highest expression level, whereas Q5 corresponds to genes with the lowest expression. The metaplot depicts the H1-3XFLAG enrichment pattern for each expression group across the TSS or TTS. (C) Heatmaps show the distribution of H1-3XFLAG across all *N. crassa* genes centered on the TSS (left) or TTS (right). Genes are ordered by expression level from highest to lowest. (D) Heatmaps show the distribution of RNA polymerase II across all *N. crassa* genes centered on the TSS (left) or TTS (right). Genes are ordered as in C. Data in B-D are from replicate one.

On average, H1-3XFLAG was enriched upstream of the TSS and was depleted from gene bodies. Similar enrichment patterns were observed for both biological replicates, but not in a negative control experiment in which we used anti-FLAG antibodies to perform ChIP-seq in a wildtype strain (no FLAG-tagged protein). We next asked if H1-3XFLAG enrichment was correlated with the level of transcription. We binned Neurospora genes into five groups based on expression level and constructed metaplots to visualize the average H1-3XFLAG distribution pattern for each group. The level of enrichment in gene promoters was positively correlated with the level of transcription (Figure 3B). Similarly, depletion of H1-FLAG from coding regions was correlated with the level of transcription. To explore this further, we performed ChIP-seq for RNA polymerase II and constructed heatmaps of H1-FLAG and RNA polymerase II occupancy across all Neurospora genes ordered by expression level (Figure 3C,D). These data further support the idea that H1-3XFLAG enrichment is highest in the promoters and lowest in the coding sequences of highly expressed genes.

### H1 is not a major determinant of nucleosome positioning in *N. crassa*

Given that H1-3XFLAG displayed a subtle preference for promoters of actively expressed genes, we hypothesized that H1 may impact chromatin structure at promoters. Like many eukaryotes, *N. crassa* promoters are characterized by nucleosome free regions (NFRs) upstream of the transcription start site (TSS) followed by well positioned “+1” nucleosomes (Sancar *et al.* 2015). To determine if H1 is important for establishment of this characteristic promoter structure and to determine if *N. crassa* H1 is important for nucleo-some occupancy or nuclesome positioning at specific sites in the genome, we performed Micrococcal nuclease digestion followed by sequencing (MNase-seq). Nuclei were incubated with MNase for increasing times (up to 60 minutes) before the DNA from each digestion reaction was purified and resolved on an agarose gel. In agreement with previously published work (Folco *et al.* 2003), wild-type and Δ*hH1* showed similar global MNase sensitivity across multiple experiments (representative gels are shown in Figure 4A and B). We next performed gel extractions to isolate the DNA band corresponding to mononucleosomes (~150 base pairs) from samples digested with MNase for 20 or 60 minutes and constructed Illumina sequencing libraries. For each digestion time, libraries were prepared from two independent biological replicates (four samples total for each strain) and paired-end sequencing was performed. We observed no differences in the average length of the mononucleosomal fragments in wildtype and Δ*hH1* (Table 1). As expected, longer digestion times produced shorter DNA fragments (compare 20 minute to 60 minute digestion times).

**Figure 4.**
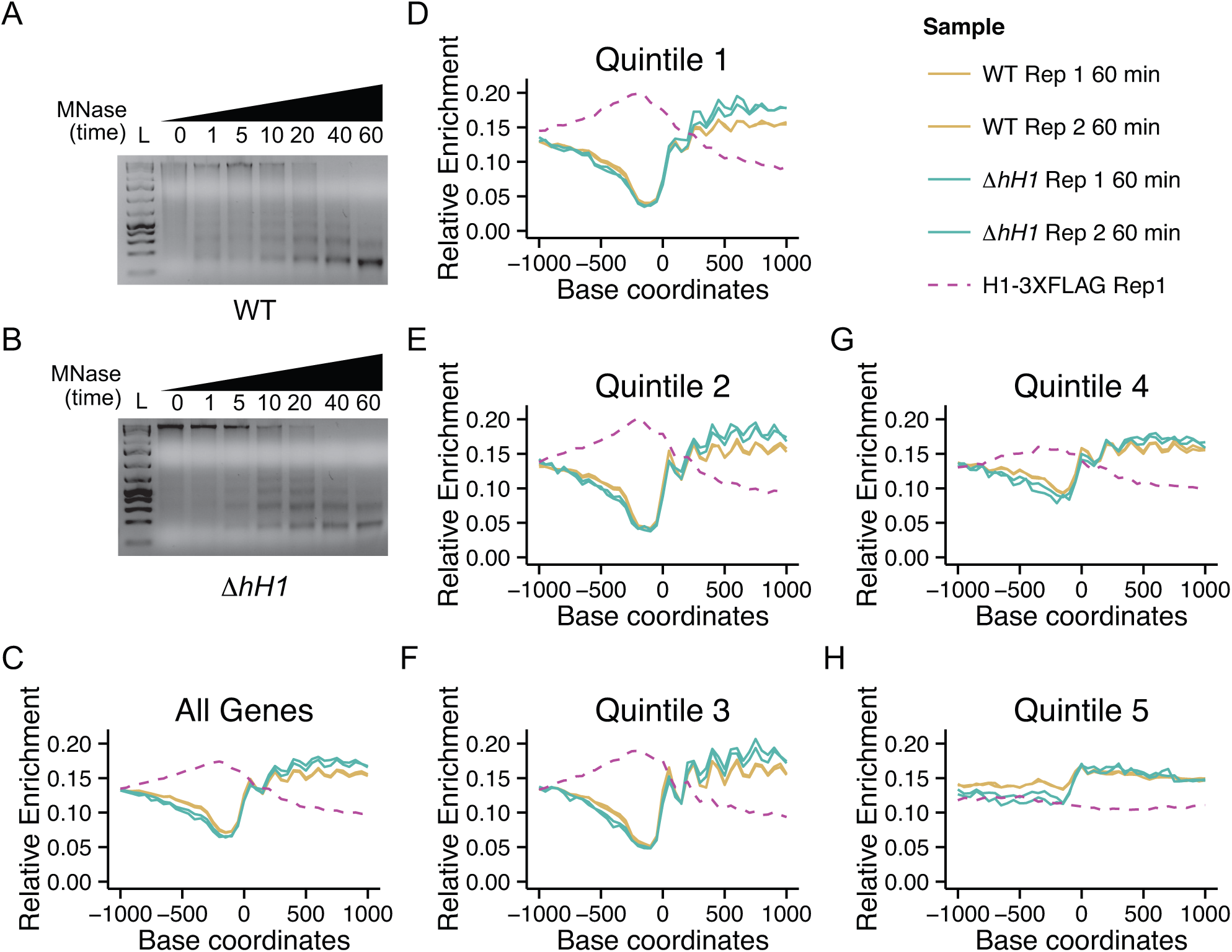
H1 is not a major determinant of nucleosome size or positioning in Neurospora. (A) Wildtype or (B) Δ*hH1* nuclei were isolated and treated with Micrococcal nuclease (MNase) for the indicated times (minutes). DNA was purified, resolved on an agarose gel, and visualized by staining with Ethidium Bromide. Wildtype and Δ*hH1* display similar digestion kinetics. (C) Metaplots depict the average sequencing depth on the + strand across all *N. crassa* genes obtained by sequencing mononucleosomal DNA from a 60 minute MNase digest. Plots show data from two biological replicates each for wildtype and Δ*hH1*. (D - H) All *N. crassa* genes were ranked by expression level and split into quintile groups, ranging from highest expression (Quintile 1) to lowest expression (Quintile 5). Metaplots depict the average sequencing coverage on the + strand for each group. Plots show data from two biological replicates each for wildtype and Δ*hH1*. The legend in the top right panel corresponds to plots (C - H). For all plots, the coverage of + strand reads from an H1-3XFLAG ChIP-seq experiment is shown as a dashed line.

**Table 1.** Paired-end read insert size

| Sample | Digest Time | Mean Insert Size <sup>a</sup> |
| --- | --- | --- |
| Wildtype replicate 1 | 20 min | 139.9 ±23.7 |
| Wildtype replicate 2 | 20 min | 144.7 ±25.1 |
| Δ*hH1* replicate 1 | 20 min | 144.1 ±22.4 |
| Δ*hH1* replicate 2 | 20 min | 139.6 ±22.7 |
| Wildtype replicate 1 | 60 min | 127.8 ±33.0 |
| Wildtype replicate 2 | 60 min | 126.5 ±28.3 |
| Δ*hH1* replicate 1 | 60 min | 129.8 ±33.9 |
| Δ*hH1* replicate 2 | 60 min | 125.6 ±24.3 |

<sup>a</sup> ±standard deviation

We next asked if H1 impacted the occupancy and/or positioning of nucleosomes in promoters or gene coding sequences. We created metaplots to analyze the average nucleosome distribution across all *N. crassa* genes (Figure 4C). In this case, only plus strand reads were plotted to generate peaks corresponding to the edge of each nucleosome. The average gene profile revealed a characteristic nucleosome free region upstream of the TSS, as reported previously for *N. crassa* (Sancar *et al.* 2015). The average nucleosome position profiles were similar in wildtype and Δ*hH1*. Because H1 was most enriched in the promoters of highly expressed genes, we next plotted average MNase-seq enrichment profiles for genes grouped by expression level. Although we detected a subtle increase in the size of the NFR from the most expressed genes in the two 60 minute digestion samples, this difference was not ap-parent in the samples subjected to a 20 minute MNase digestion (**Figure S1**). The difference was also not apparent in a third independent replicate subjected to single-end Illumina sequencing (**Figure S1**). Thus, we conclude that H1 is not a major determinant of nucleosome positioning or occupancy in Neurospora. Because it appeared that enrichment of H1-3XFLAG was inversely correlated with nucleosome occupancy, we plotted plus strand reads from replicate one of the H1-3XFLAG ChIP-seq experiment described above to allow direct comparison of nucleosome occupancy and H1-3XFLAG enrichment (Figure 4C - H). This confirmed that the highest levels of H1 enrichment correspond to sites with the lowest nucleosome occupancy, raising the possibility that H1-3XFLAG binds to DNA that is free of nucleosomes as well as to nucleosomal DNA in *N. crassa*.

### Δ*hH1* exhibits increased DNA methylation in *N. crassa* heterochromatin domains

H1 impacts DNA methylation levels in animals (Fan *et al.* 2005a; Zhang *et al.* 2012a), plants (Wierzbicki and Jerzmanowski 2005; Zemach *et al.* 2013; Rea *et al.* 2012), and the fungus *Ascobolus im-mersus* (Barra *et al.* 2000). It was previously reported that H1 did not impact DNA methylation in *N. crassa* (Folco *et al.* 2003); however, Folco and colleagues analyzed only a single methylated region. It remained possible that H1 impacts DNA methylation in a region-specific manner or alternatively, that H1 has subtle impact on DNA methylation that may have been overlooked. We performed MethylC-seq to analyze DNA methylation across the entire genome at single base-pair resolution. Genomic DNA was isolated from two replicates each of a wildtype *mat A* strain, the Δ*hH1* mutant, and a negative control Δ*dim-2* strain, which lacks DNA methylation altogether (Kouzminova and Selker 2001). All strains were grown simultaneously, but the wildtype and Δ*dim-2* data were published previously as part of another study (Basenko *et al.* 2016).

To determine if H1 impacts the level or distribution of DNA methylation in *N. crassa*, we first plotted DNA methylation levels across Linkage Group VII, a 4Mb chromosome corresponding to ~10% of the genome. The overall pattern of DNA methylation was similar in wildtype and the Δ*hH1* strain. However, we noted that DNA methylation levels were higher in the Δ*hH1* mutant at most regions along the chromosome (Figure 5A). We next constructed metaplots to quantify the average methylation level for all genomic regions that are normally methylated in the wildtype *mat A* strain (see Materials and Methods). Both Δ*hH1* replicates displayed higher average DNA methylation levels when compared to the wildtype strain (Figure 5B). To confirm that this wasn’t an artifact due to differences in strain backgrounds, we compared the level of methylation in the Δ*hH1* strain to a wildtype mat a strain and we analyzed DNA methylation levels in a second hH1 loss of function strain in which the hH1 gene was inactivated by repeat-induced point mutation (Folco *et al.* 2003). In both cases, the level of DNA methylation was higher in the *hH1* mutant strains (Figure 5C,D). A search for differentially methylated regions between wildtype and Δ*H1* identified only a single methylated region that was specific to the Δ*H1* strain (Linkage Group VI; 309590 - 319133). This hyper-methylated sequence corresponds to *Sly1-1*, a DNA transposon found in the genomes of some *N. crassa* isolates (Wang *et al.* 2015).

**Figure 5.**
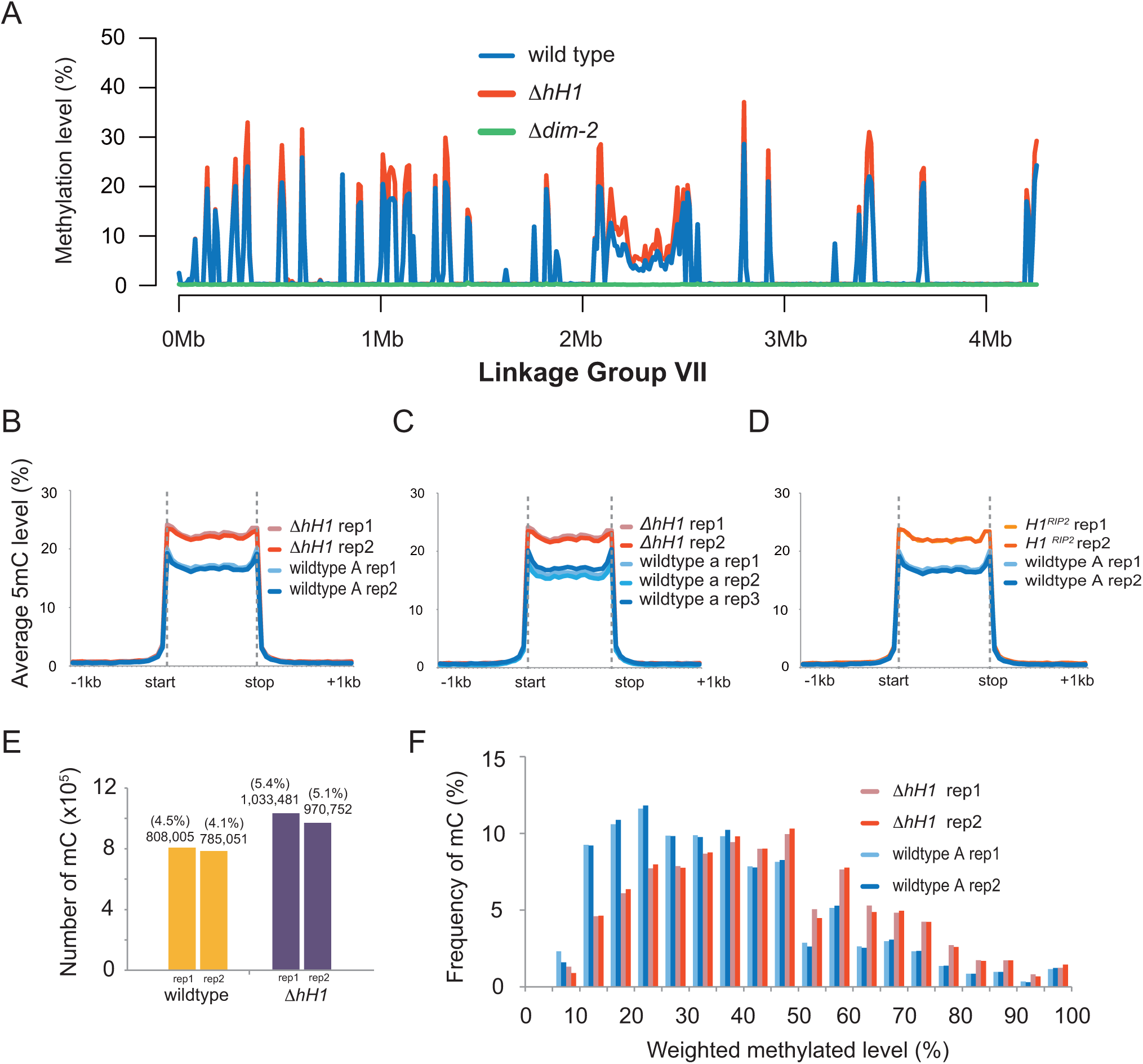
Increased DNA methylation is observed at most *N. crassa* heterochromatin domains in H1-deficient strains. (A) The DNA methylation level (weighted DNA methylation level (%); see Materials and Methods) is shown for 10 Kb windows across *N. crassa* Linkage Group VII for wildtype, the Δ*hH1* mutant, and the Δ*dim-2* strain, which lacks all DNA methylation. (B-D) The metaplots show the average DNA methylation level across all previously identified wildtype methylated domains: (B) the wildtype *mat A* strain and the Δ*hH1* strain; (C) the wildtype mat a strain and the Δ*hH1* strain; and (D) the wildtype *mat A* strain and the *hH1*^*RIP2*^ strain. Data for at least two independent biological replicates of each strain are shown. (E) More cytosines are methylated in Δ*hH1*. The plot shows the total number of methylated cytosines (see Materials and Methods) identified in each wildtype and Δ*hH1* replicate. (F) The level of methylation at individual cytosines is higher in ΔhH1. The percentage of total shared methylated sites (y-axis) versus the level of methylation at individual cytosines (x-axis) is shown.

Higher methylation levels can result when individual cytosines are methylated at a higher frequency in a population of nuclei or when new cytosines become methylated (or a combination of both). To determine which of these possibilities was the case in H1-deficient strains, we determined the number of methylated cytosines in wildtype and Δ*hH1* using a binomial test in combination with multiple testing correction (see Materials and Methods). The number of methylated cytosines increased by ~25% in the Δ*H1* strain. We next compared methylation frequency at cytosines that were scored as methylated in both strains. This revealed that shared sites are methylated at higher frequency in nuclei from the Δ*hH1* strain. Thus, loss of H1 leads to a subtle, but global increase in DNA methylation at relics of repeat-induced point mutation.

## DISCUSSION

We applied genomic and molecular methods to investigate H1 in the model fungus *N. crassa*. We first confirmed that Neurospora H1 is a chromatin component by constructing a functional epitope-tagged H1 fusion protein and analyzing its localization in cell fractionation and ChIP-seq experiments. This revealed that H1 is globally enriched throughout the genome, occupying both heterochromatic and euchromatic regions. Surprisingly, we observed the highest enrichment of H1-3XFLAG in promoters of expressed genes. Although the overall amplitude of enrichment was low, preferential enrichment of H1-3XFLAG was clearly correlated with expression level. These results may indicate that H1 plays different roles in fungi and animals. In mouse embryonic stem cells (ESCs), H1c and H1d are depleted from strong promoters and enhancers (Cao *et al.* 2013). It should be noted that H1 protein levels are significantly reduced in ESCs compared to differentiated cells (Fan *et al.* 2003), which might explain why H1c and H1d are not detected at ESC promoters. This seems unlikely, however, as ChIP-seq analyis of H1b from ESCs and from differentiated cells revealed that this protein was similarly enriched at repressed genes and depleted from active promoters (Li *et al.* 2012a). On the other hand, given that mammals encode multiple H1 variants it is possible that certain H1 variants will bind preferentially to nucleosome-depleted promoter regions, similar to the case for Neurospora H1. Indeed, analysis of H1 variants by DamID showed that H1.1 exhibits a distinct localization compared to other histones and is not excluded from promoters like the other somatic H1 variants (Izzo *et al.* 2013). Similarly, the more divergent H1.X variant was recently found to be enriched at regions with high RNA polymerase II occupancy (Mayor *et al.* 2015), suggesting that these H1 variants may play roles within active chromatin in higher eukaryotes. Additional work to define the specific roles of linker histones in fungal and animals systems is needed to determine their specific functions at open chromatin.

One possibility is that *N. crassa* H1 functions more like animal HMG (High Mobility group) proteins. The chromatin architectural protein HMGD1 is enriched in active chromatin, for example (Nalabothula *et al.* 2014), similar to the results obtained for H1-3XFLAG here. It is important to note that we can not rule out the possibility that the subtle promoter enrichment observed for H1-FLAG is an experimental artifact. This could be the case if the basic H1 protein interacts non-specifically with DNA sequences originating from open chromatin during the ChIP procedure. It was shown that certain highly expressed loci in yeast are prone to artifactual enrichment in ChIP-chip experiments for reasons that are not understood (Teytelman *et al.* 2013). However, we do not think this is the case here because the non-specific enrichment patterns observed in yeast are qualitatively different than the enrichment patterns we observe for H1-3XFLAG. Thus, it is likely that the patterns observed here reflect the in vivo occupancy of H1. It will be interesting to determine if promoter regions interact with a sub-population of H1 in which specific residues are post-translationally modified. It will also be interesting to determine if depletion of H1 from coding sequences of expressed genes depends on post-translational modification. This seems likely given that H1 proteins in plants and animals are extensively modified, much like the core histones (Harshman *et al.* 2013; Bednar *et al.* 2015; Annalisa and Robert 2015; Kotliński *et al.* 2016). Moreover, it was shown that phosphorylation of H1 was linked to transcription by RNA polymerase I and II in humans (Zheng *et al.* 2010). An important goal for future studies will be to determine if *N. crassa* H1 is post-translationally modified and to determine if different forms of H1 exhibit distinct localization and/or distinct functions.

We found here that deletion of *hH1* from *N. crassa* did not substantially alter global MNase accessibility or nucleosome positioning. Moreover, H1 did not impact the size of protected DNA fragments produced by MNase treatment or the distance between adjacent nucleosomes in genes. These results point to clear differences in how H1 interacts with chromatin in *N. crassa* and in animals. Indeed, H1 depletion caused increased MNase accessibility, altered nucleosome spacing lengths, and reduced chromatin compaction in H1 triple-knockout ESCs (Fan *et al.* 2005b). Our results could indicate that *N. crassa* H1 does not bind to the linker DNA and the dyad axis of the NCP as demonstrated for animal H1 (Bednar *et al.* 2015). Another possibility is that *N. crassa* H1 is more dynamic than H1 in higher eukaryotes. FRAP studies revealed that mammalian H1 variants exist in high-mobility and low-mobility pools and that the half-life of fluorescence recovery after H1-GFP bleaching was significantly shorter than for the core histones (Misteli *et al.* 2000; Lever *et al.* 2000). Interactions between H1 and the NCP may be even more transient in *N. crassa* such that H1 does not interfere with MNase digestion even though it interacts with the same region of the nucleosome protected by animal H1.

We found increased DNA methylation in H1-deficient cells of *N. crassa*. H1 effects DNA methylation in both *A. thaliana* and animal cells, but the relationship between H1 and DNA methylation is different in these systems. In animals, H1 variants promote repressive modifications, including DNA methylation in mammals (Yang *et al.* 2013) and H3K9me2 in both mammals and Drosophila (Li *et al.* 2012a; Lu *et al.* 2013). The observation that H1-deficient cells exhibit hypermethylation demonstrates that *N. crassa* H1 is not required to promote DNA methylation or H3K9 methylation, which directs DNA methylation in Neurospora (Tamaru and Selker 2001). Similar hypermethyaltion was reported in *A. immersus* (Barra *et al.* 2000). Moreover, the DNA methylation phenotypes of *N. crassa* and *A. immersus* are reminiscent of *A. thaliana* H1 depletion lines, where a global increase in DNA methylation was observed in heterochromatin domains along with loss of DNA methyltion in euchromatic transposon sequences (Wierzbicki and Jerzmanowski 2005; Zemach *et al.* 2013). These results indicate that H1 can limit DNA methylation in plants and fungi. Indeed, depletion of *A. thaliana* H1 rescued the reduced DNA methylation phenotype of ddm1 plants, leading to the conclusion that DDM1 promotes DNA methyaltion by removal of H1. This may indicate that H1 evolved a new function to promote heterochromatic modifications in the animal lineage.

It is possible that *N. crassa* H1 limits access of the DNA methyl-transferase DIM-2 or the H3K9 MTase DIM-5^KMT1^. A recent study showed that binding of H1 to the NCP limited the dynamics and modifiability of the H3 tail in vitro (Stützer *et al.* 2016), consistent with this possibility. In addition, increased accessibility of the DIM-2 DNA methyltransferase was linked to hypermethylation in the *N. crassa* histone deacetylase-1-deficient strain (Honda *et al.* 2012). On the other hand, in *A. thaliana* H1 was required for imprinting of the MEDEA locus, which involves active removal of methyl cytosine bases by DNA glycosylases (Rea *et al.* 2012). A mechanism for DNA demethylation has not been described in *N. crassa*, but it is possible that H1 promotes removal of methylated cytosines in heterochromatin domains. Additional studies are needed to understand exactly how H1 impacts DNA methylation levels in *N. crassa*. Overall, this work adds to the diverse set of phenotypes that have been reported following depletion of H1 in plants, animals, and fungi. Future work to investigate H1 in fungal systems is likely to yield new insights into the evolution and the functions of this important group of proteins.

## ACKNOWLEDGMENTS

We would like to thank Cameron Prybol for technical contributions to the project. This work was funded by a grant from the American Cancer Society to ZAL (RSG-14-184-01-DMC), a grant from the National Institutes of Health to RJS (R00GM100000), and a grant to XZ from the National Science Foundation (#0960425).

